# Genomic analyses identify key molecules and significant signaling pathways in Sorafenib treated hepatocellular cancer cells

**DOI:** 10.1101/2022.03.15.484435

**Authors:** Tingxiang Chang, Mengshan Wang

## Abstract

Sorafenib (Sfb) is a multikinase inhibitor that is significantly effective in killing liver cancer. However, the potential signaling pathways and mechanisms remain unknown. Here, we aim to identify the functional molecules and signaling pathways by analyzing the RNA-seq data. The GSE186280 was produced by the Illumina HiSeq 2000 (Homo sapiens). The KEGG and GO analyses indicated the MAPK signaling pathway and transcriptional misregulation in cancer are the major signaling pathways during the treatment of Sfb in liver cancer. Moreover, we identified ten key molecules including TP53, MYC, ACTB, CREBBP, HSP90AA1, JUN, UBB, SIRT1, CCNA2, RBX1. Therefore, this study provides novel knowledge of Sfb treated liver cancer.

## Introduction

Liver cancer is a developing health challenge worldwide. By 2025, more than one million population will be impacted by cancer annually^1^. Hepatocellular carcinoma (HCC) is the most common form of liver cancer and accounts for nearly 90% of cases^2^. Hepatitis B virus (HBV) is the primary reason for liver cancer^3^. Non-alcoholic steatohepatitis (NASH), associated with metabolic syndrome is becoming the fastest growing etiology of HCC^4^. About 25% of HCC cancers present actionable mutations^5^. Indeed, the most common mutational drivers in HCC, such as TERT and TP53 remain undruggable^6^. Additionally, translational therapies are still under investigation. Currently, targeting the tumor microenvironment has provided new insights into drug development^7^. The diagnosis of HCC is largely based on the non-invasive criteria but there is a growing need for confirmation by using tissue biopsies in clinical practice^8^.

The novel therapies including immune-checkpoint inhibitors and monoclonal antibodies have challenged the application of conventional therapies^9^. Most patients with liver cancer are estimated to be continuously using systematic therapies in their life span^10^. By application of systemic therapies in the past 5 years, the overall survival and the quality of patients have markedly increased^11^.

In this study, we explored the effects of using Sorafenib (Sfb) on liver cancer by using the RNA-seq data. We identified several DEGs and significant pathways through the KEGG and GO. We also created the Reactome map and protein-protein interaction (PPI) network to study the relationships among the DEGs. These DEGs and biological processes will provide guidance on the treatment of liver cancers.

## Methods

### Data resources

Gene dataset GSE186280 was obtained from the GEO database. The data was produced by the Illumina HiSeq 2000 (Homo sapiens) (Synthesis and functions of ribosomes, University of Seville, Antonio Maura Montaner s/n. IBiS, Seville, Seville). The analyzed dataset includes 3 groups of controls and 3 groups of Sorafenib (Sfb) treated Hepatocellular carcinoma (HCC) cells.

### Data acquisition and processing

The data were organized and conducted by the R package as previously described^12–15^. We used a classical t-test to identify DEGs with P< 0.01 and fold change ≥1.5 as being statistically significant.

The Kyoto Encyclopedia of Genes and Genomes (KEGG) and Gene Ontology (GO) KEGG and GO analyses were conducted by the R package (ClusterProfiler) and Reactome^16–19^. P<0.05 was considered statistically significant.

### Protein-protein interaction (PPI) networks

The Molecular Complex Detection (MCODE) was used to construct the PPI networks. The significant modules were produced from constructed PPI networks^20–23^. The pathway analyses were performed by using Reactome (https://reactome.org/), and P<0.05 was considered significant.

## Results

### Identification of DEGs in hepatocellular cancer cells with the treatment of Sfb

To determine the impact of Sfb on hepatocellular cancer cells, we analyzed the RNA-seq data from HepG2 cells with the treatment of Sfb. A total of 737 genes were identified with the threshold of P < 0.01. The up-and-down-regulated genes were shown by the heatmap and volcano plot (Figure 1). The top ten DEGs were selected in Table 1.

**Figure 1.**
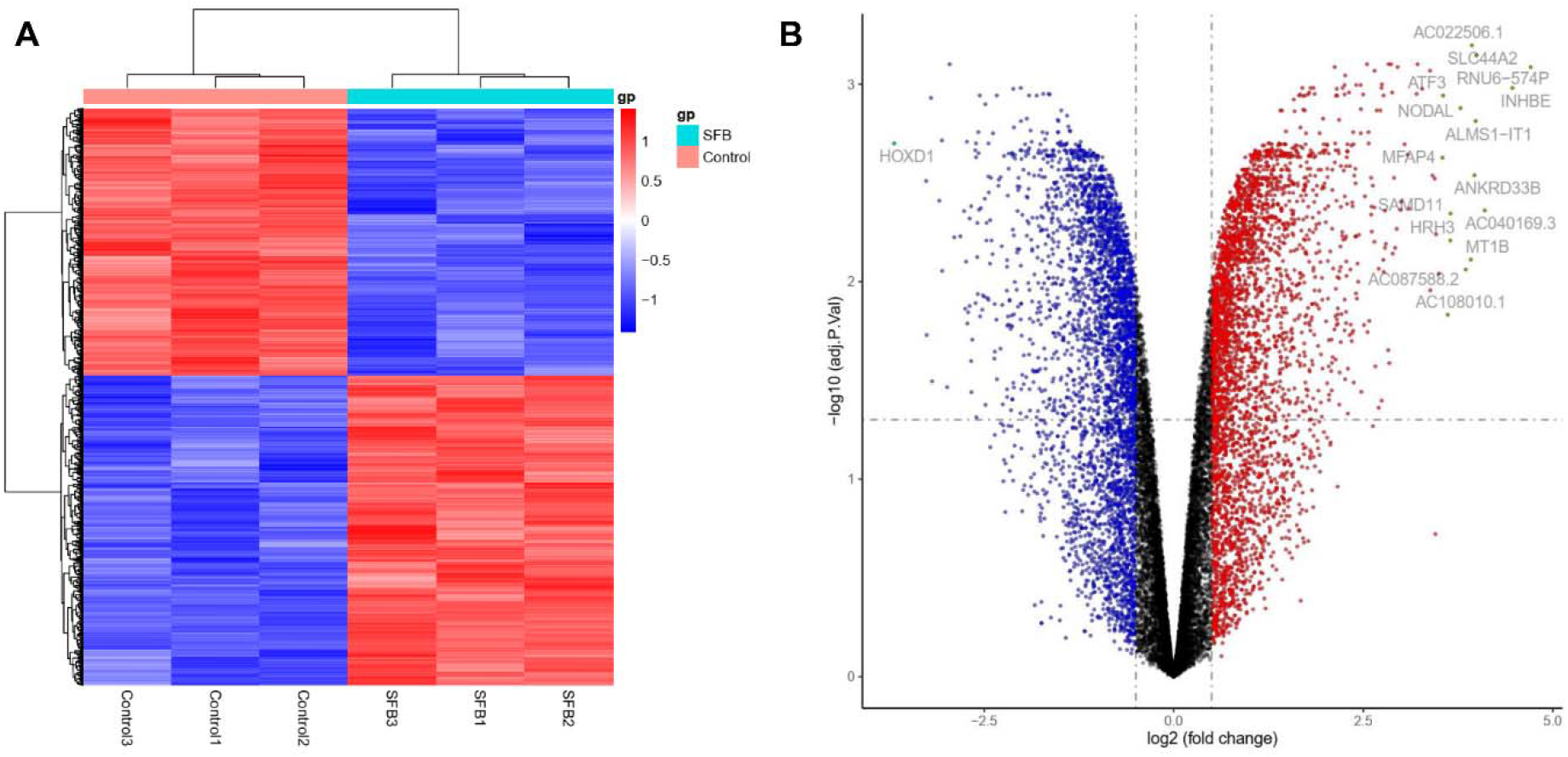
Heatmap and volcano plot were created in hepatocellular cancer cells with the treatment of Sfb. (A) Significant DEGs (P < 0.01) were used to produce the heatmap. (B) Volcano plot for DEGs in hepatocellular cancer cells with the treatment of Sfb. The most significantly changed genes are highlighted by grey dots.

**Table 1.**

| **Entrez gene** | **Gene symbol** | **Fold-change** | **Regulation** |
| --- | --- | --- | --- |
| Top 10 down-regulated DEGs |  |  |  |
| 3231 | HOXD1 | -3.687511531 | Down |
| 100216001 | MANCR | -3.26533599 | Down |
| 3303 | HSPA1A | -3.202987082 | Down |
| N/A | RF02160 | -3.091519071 | Down |
| 100507410 | C1QTNF1-AS1 | -3.058048145 | Down |
| N/A | AC015813.5 | -2.956365835 | Down |
| 147495 | APCDD1 | -2.765486709 | Down |
| 100506343 | INKA2-AS1 | -2.729246819 | Down |
| 100271234 | RPS20P21 | -2.692314243 | Down |
| 100128191 | TMPO-AS1 | -2.671939984 | Down |
| Top 10 up-regulated DEGs |  |  |  |
| 106479820 | RNU6-574P | 4.707285027 | up |
| 83729 | INHBE | 4.467070002 | up |
| N/A | AC040169.3 | 4.102029975 | up |
| 57153 | SLC44A2 | 3.990570932 | up |
| 100874291 | ALMS1-IT1 | 3.982717054 | up |
| 651746 | ANKRD33B | 3.966034948 | up |
| N/A | AC022506.1 | 3.931997903 | up |
| 4838 | NODAL | 3.784543224 | up |
| 148398 | SAMD11 | 3.651319779 | up |
| 467 | ATF3 | 3.552812261 | up |

### Identification of KEGG and GO in hepatocellular cancer cells with the treatment of Sfb

To further determine the functions of Sfb in hepatocellular cancer cells, we applied the KEGG and GO analysis (Figure 2). We identified the top ten KEGG pathways including “MAPK signaling pathway”, “Transcriptional misregulation in cancer”, “Signaling pathways regulating pluripotency of stem cells”, “TNF signaling pathway”, “Apoptosis”, “Colorectal cancer”, “p53 signaling pathway”, “Steroid biosynthesis”, “Terpenoid backbone biosynthesis”, and “Mismatch repair”. We identified the top ten biological processes of GO, including “Intracellular receptor signaling pathway”, “Cellular response to transforming growth factor beta stimulus”, “Response to transforming growth factor beta”, “Transforming growth factor beta receptor signaling pathway”, “Regulation of DNA-templated transcription in response to stress”, “Sterol biosynthetic process”, “Regulation of transcription from RNA polymerase II promoter in response to stress”, “Cholesterol biosynthetic process”, “Secondary alcohol biosynthetic process”, “Positive regulation of transcription from RNA polymerase II promoter in response to stress”. We identified the top ten cellular components of GO, including “Nuclear speck”, “Transcription regulator complex”, “RNA polymerase II transcription regulator complex”, “Protein-DNA complex”, “Histone acetyltransferase complex”, “Protein acetyltransferase complex”, “Acetyltransferase complex”, “H4 histone acetyltransferase complex”, “mRNA cleavage factor complex”, and “DNA replication preinitiation complex”. We then identified the top ten molecular functions, including “Transcription coregulator activity”, “DNA-binding transcription factor binding”, “DNA-binding transcription activator activity, RNA polymerase II-specific”, “Protein serine/threonine kinase activity”, “RNA polymerase II-specific DNA-binding transcription factor binding”, “Magnesium ion binding”, “Transcription corepressor activity”, “Histone deacetylase binding”, “SMAD binding”, and “RNA methyltransferase activity”.

**Figure 2.**
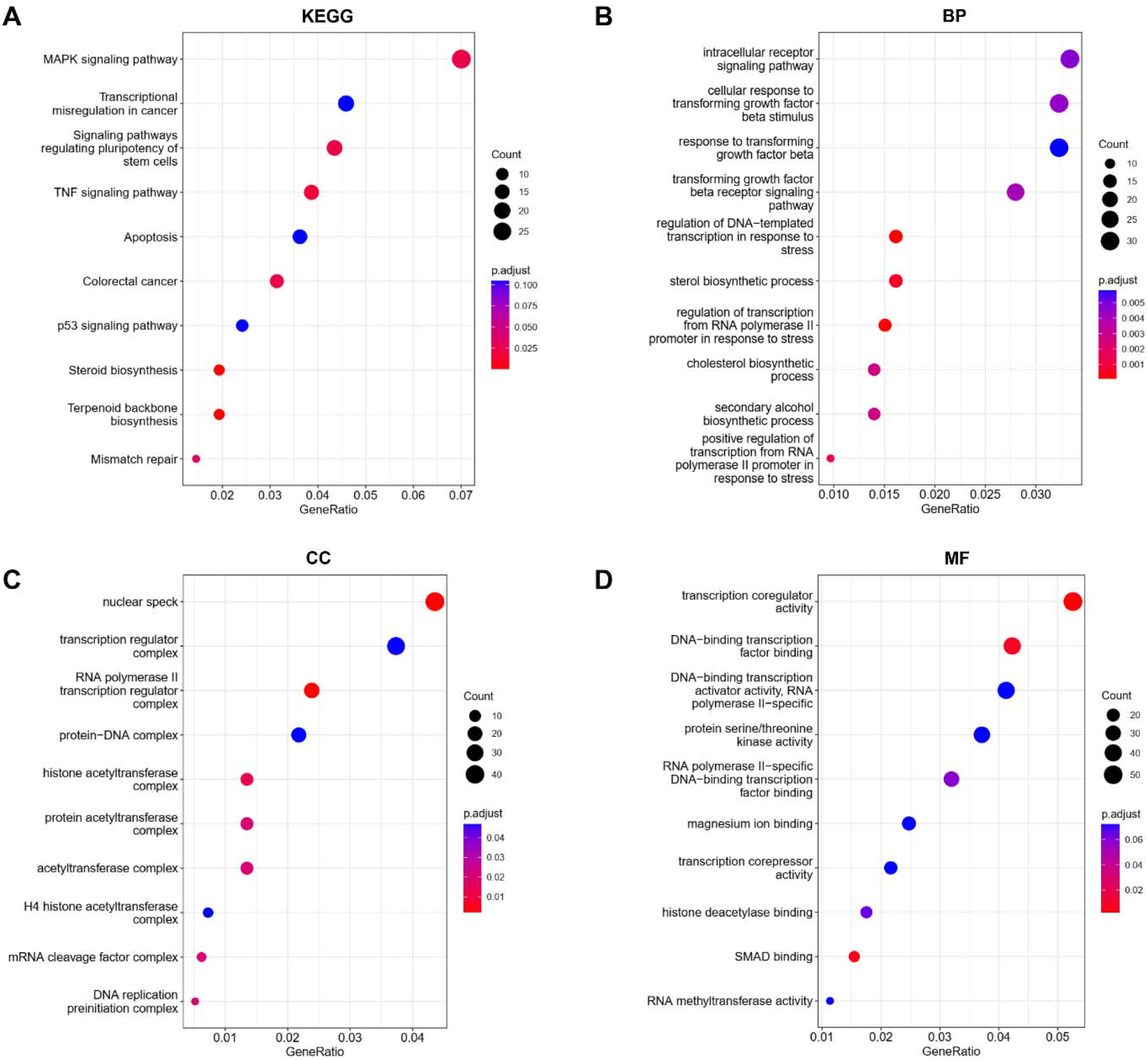
KEGG and GO analyses of DEGs in hepatocellular cancer cells with the treatment of Sfb. (A) KEGG analysis, (B) Biological processes, (C) Cellular components, (D) Molecular functions.

### PPI networks in hepatocellular cancer cells with the treatment of Sfb

To determine the interaction among the DEGs, we constructed the PPI networks by using the Cytoscape software (combined score > 0.4). Table 2 showed the top ten interactive molecules with the highest degree scores. The top two modules were shown in Figure 3. We further examined the mechanisms with Reactome map (Figure 4) and identified the top ten significant processes including “Activation of gene expression by SREBF (SREBP)”, “Regulation of cholesterol biosynthesis by SREBP (SREBF)”, “Response of EIF2AK1 (HRI) to heme deficiency”, “Attenuation phase”, “HSF1-dependent transactivation”, “HSF1 activation”, “ATF4 activates genes in response to endoplasmic reticulum stress”, “TP53 Regulates Transcription of Genes Involved in Cytochrome C Release”, “Activation of PUMA and translocation to mitochondria”, and “Activation of BH3-only proteins” (Supplemental Table S1).

**Figure 3.**
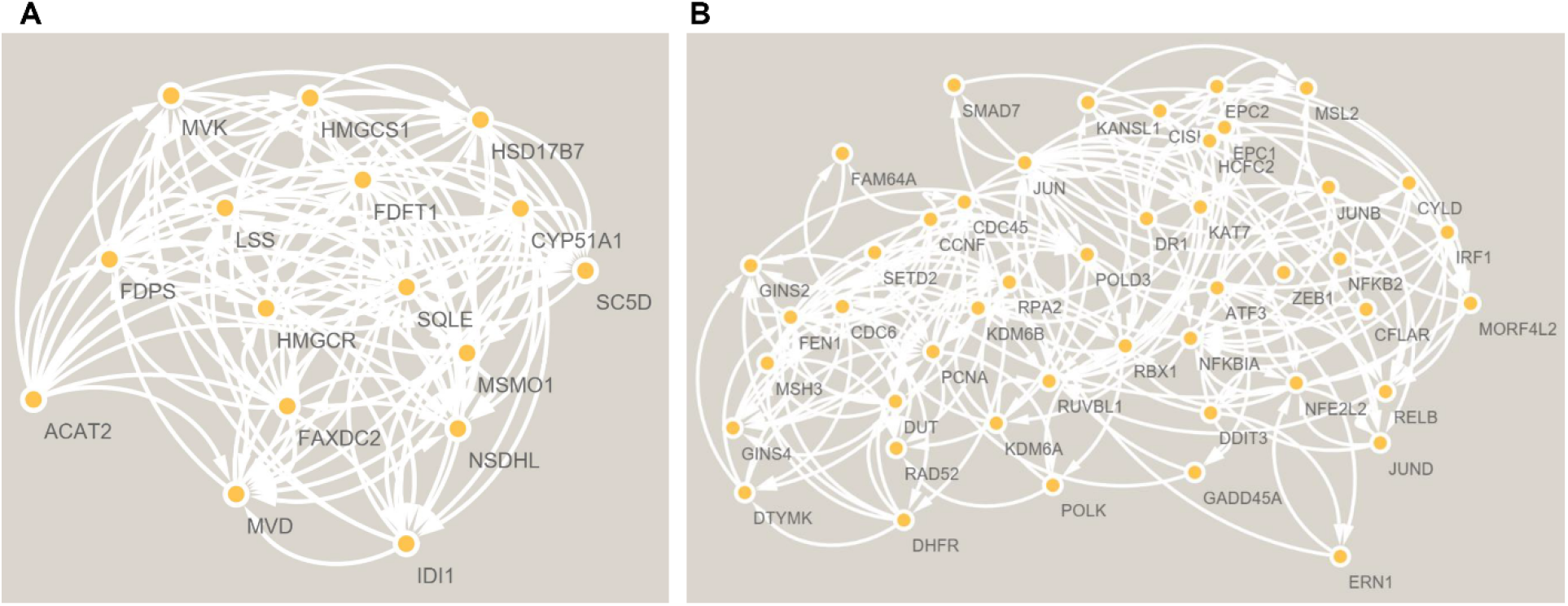
The PPI network analyses of DEGs in hepatocellular cancer cells with the treatment of Sfb. The cluster (A) and cluster (B) were constructed by MCODE.

**Figure 4.**
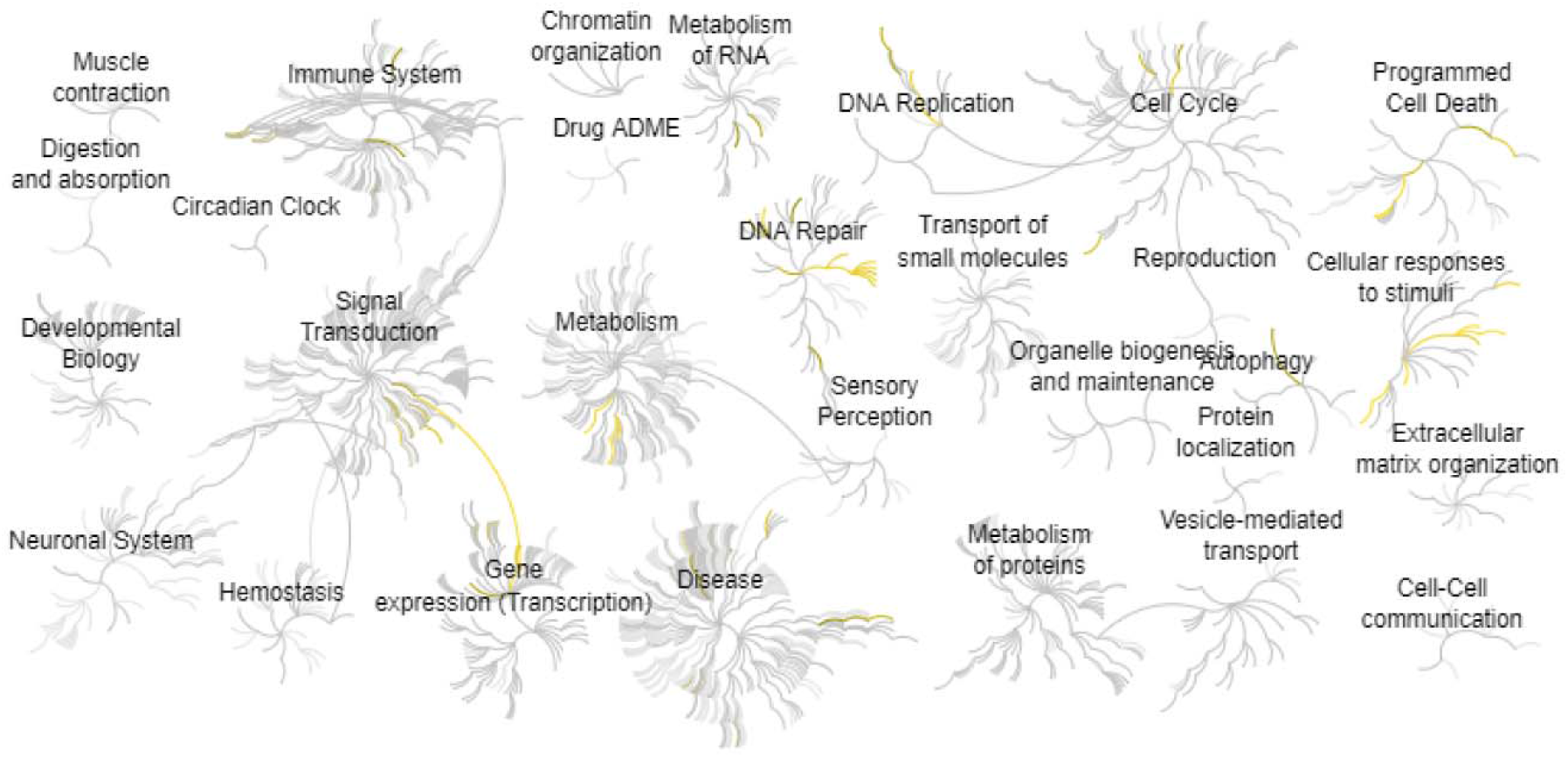
Reactome map representation of the significant biological processes in hepatocellular cancer cells with the treatment of Sfb

**Table 2.** Top ten genes demonstrated by connectivity degree in the PPI network.

| **Gene symbol** | **Gene title** | **Degree** |
| --- | --- | --- |
| TP53 | Tumor protein p53 | 155 |
| MYC | MYC proto-oncogene, bhlh transcription factor | 132 |
| ACTB | Actin beta | 106 |
| CREBBP | CREB binding protein | 86 |
| HSP90AA1 | Heat shock protein 90 alpha family class A member 1 | 83 |
| JUN | Jun proto-oncogene, AP-1 transcription factor subunit | 80 |
| UBB | Ubiquitin B | 80 |
| SIRT1 | Sirtuin 1 | 64 |
| CCNA2 | Cyclin A2 | 60 |
| RBX1 | Ring-box 1 | 50 |

## Discussion

Liver cancer is the second leading cause of cancer-related mortality in developing countries^1^. The proposed “precision medicine for liver cancer” holds hopes for novel therapy of liver cancer, which was designed to help doctors choose treatments based on the heterogeneity of patients^24^. Thus, our study further explored the mechanism of Sfb treated liver cancer cells.

We found the MAPK signaling pathway and transcriptional misregulation in cancer are the major signaling pathways according to KEGG and GO analyses. The study by Bénédicte Delire et al showed that the MAPK signaling pathway and their effectors should be considered potential therapeutic targets in the treatment of liver cancer. Especially, after the treatment of Srb, more MAPK repressors have merged and are under investigation for liver cancer therapy^25^. Similarly, Hyuk Moon found that MAPK is frequently activated in liver cancer, which follows by a number of genetic mutations in Hepatocellular cancer cells^26^. The study by Yin Yuan et al. found that lincZNF337 can acetylate the H2A via KAT5 to further transcriptionally misregulate the cancer signaling pathway in liver cancer^27^.

Besides the analysis of signaling pathways, we also identified several interactive proteins. Galina Khemlina et al found that TP53 is one of the most common molecular anomalies in liver cancer^28^. MYC has associated with the aurora kinase A protein in a complex that acts as a drug target in P53-altered liver cancer^29^. Circadian gene clocks and their regulators are involved in different signaling pathways including inflammation, metabolism, aging, and cancer^30^. Interestingly, MYC can control the circadian clocks and metabolism in cancer cells^31–40^. Madeleine E Lemieux et al demonstrated the inactivation of Crebbp selectively alters mRNA processing in hematopoietic stem cells, which can cause gene mutation and cancer genesis ^41^. Weidong Shi et al found that FBXL6 controls the c-MYC to drive the progression of liver cancer through ubiquitination and stabilization of HSP90AA1^42^. Yan Chen et al found the O-GlcNAcylated c-Jun represses the ferroptosis through inhibiting GSH synthesis in liver cancer^43^. Li Nan et al found UBB is an important liver cancer marker that is increased in the cancer progression ^44^. SIRT1 is increased in cancer cells that may play a critical role in tumor initiation, progression, and angiogenesis^45^. GPCR proteins and their regulators control numerous signaling pathways and biological processes such as inflammation, cancer, aging, and metabolism^46–58^. Gα12 can regulate SIRT1 protein through the HIF-1α-mediated transcriptional induction of ubiquitin-specific peptidase 22^59^. Lu Zeng et al found CCNA2 is a key PPI network protein that is associated with the prognosis of liver cancer patients^60^. RBX1 is one of the important targets of RhoB for degradation in liver cancer^61^.

In conclusion, our study found a key role of Sfb treatment in liver cancer. The MAPK signaling pathway and transcriptional misregulation are the major signaling pathways during Sfb treatment. Our study thus provides valuable insights into liver cancer therapy.

## Author Contributions

Tingxiang Chang: Methodology and Writing. Mengshan Wang: Conceptualization, Writing-Reviewing and Editing.

## Funding

This work was not supported by any funding.

## Declarations of interest

There is no conflict of interest to declare.

## Notes

### Competing Interest Statement

The authors have declared no competing interest.

